# WiNGS: Widely integrated NGS platform for federated genome analysis

**DOI:** 10.1101/2022.06.23.497325

**Authors:** Haleh Chizari, Nishkala Sattanathan, Amin Ardeshirdavani, Nasim Shabani, Benjamin Huremagic, Joris Robert Vermeesch, Yves Moreau, Geert Vandeweyer

## Abstract

Next-generation sequencing (NGS) has been increasingly used in a wide range of research communities and in routine clinical practice and leads to an ever increasing amount of sequencing data. Sequencing data comes with, several challenges such as sharing, storing, integrating, analyzing, and interpretion. The management of the expanding amount of data is challenging and, especially for human omics data, privacy protection is crucial. Unraveling the causes of rare diseases is critically dependent on data sharing, but progress is hampered by regulations and privacy concerns. To overcome the concerns associated with centralized human genomic data storage, we developed a federated analysis platform, referred to as Widely Integrated NGS (WiNGS). The presented approach enables datasharing and combined data-analysis of omics data across a consortium without a centralized data store. Moreover, the platform incorporates extensive variant interpretation tools from genotype to phenotype for the diagnosis of rare developmental disorders.

## 1. Introduction

Whole-exome and whole-genome sequencing (WES/WGS) data generation has exponentially increased our knowledge of the causes of rare genetic disorders[2][11]. Unraveling the genetic causes of rare diseases requires sharing of both genome and phenome information of affected individuals. In clinical diagnosis, the identification of causative variants amongst thousands or millions of candidates requires reference databases disentangling benign from potential causative variants. Hence, both the clinical interpretation of rare genetic variants and the delination of novel genetic disorders, is enhanced by the existence of large genomic and phenomic databases. Discovery of new disease genes is becoming more and more dependent on collaborative research projects, as their respective prevelances in the patient population become very small. This is usually done via centralized datasharing using high-performance infrastructure. However, the privacy sensitive nature of both genomic and clinical data make it challenging for centralized databases to conform to stringent patient privacy regulations, such as the European GDPR legislation.

Two main approaches are taken by these data sharing efforts, being either publicly sharing high-level anonymized data, or providing access to full data using restrictedaccess rights. The European Genome-phenome Archive (EGA) is a resource that facilitates access and management for long-term archival of potentially identifiable genetic, phenotypic, and clinical data in biomedical research. It provides data set discovery through publication, variant, and public meta data [12]. The 1000 Genomes Project is one of the largest distributed data collection and analysis projects [13]. The database of Genotypes and Phenotypes (dbGaP) Data Browser was developed to archive and distribute data from genotype/phenotype studies, providing individual-level datasets or summary-level information within the controlled-access portions of the database [14]. The Beacon Project is a Global Alliance for Genomics & Health (GA4GH) initiative to share both clinical and genomic data across federated networks and it provides yes/no responses about the presence of specific variants within a specific cohort. It may optionally disclose metadata, along with a “yes” response [7].

Although these platforms offer well-regulated access to sensitive data, they typically lack the full stack of interpretational and traceability methods required for clinical genomics. In the last decade, several applications have been developed to expose relevant NGS analysis tools to researchers and technicians with limited backgrounds in bioinformatics. Commercial tools, such as Golden Helix (https://www.goldenhelix.com/), genoox (https://www.genoox.com/), GeneGrid [4], or Illumina baseSpace (https://www.illumina.com/products/by-type/informatics-products/basespace-sequence-hub.html), tend to confine customers to the walled garden of the supplier, are often expensive, and can add considerable cost and maintenance overhead. Alternatively, several free or open source tools are available, including VariantDB [21], Highlander (https://sites.uclouvain.be/highlander/), VCF-server [5], VCF.filter [3], Maser [10], VarFish [6], VarAFT [8], CSI NGS Portal [9], and NGS-Logistics [1]. These tools typically provide predefined or user-based annotations and filtering strategies, but can present significant shortcomings with regard to end-to-end functionality or privacy-oriented collaboration.

For example, Highlander, VCF.Filter, and VCF-Server are deployed as standalone applications with extensive interpretational strategies, making them secure but inapplicable for collaborative projects. Although VCF-Server has a public web portal with the option to share samples amongst users, it does require central data deposition. Alternatively, Highlander enables collaborative analysis within a center by deploying a central database. Web-based analysis platforms such as CSI NGS Portal and Maser provide various fully automated pipelines to analyze NGS data and allow sharing data with consent of the data controller. However, these applications present results in static formats such as pdf reports or tab-delimited formats, making filtering less intuitive. Finally, VariantDB and VarFish are web applications for interactive variant analysis and prioritization based on pathogenicity scores and HPO terms with a focus on rare disease. When using in-house servers, these platforms allow collaborative analysis in a secure setting. Alternatively, VariantDB offers a public instance similar to VCF-Server where data, although submitted to the central database, can be shared across centers based on fine-grained access control settings. Furthermore, it offers full end-to-end API control, enabling automation in routine pipelines, from variant annotation to report generation.

With all of the above efforts aimed at one or more aspects of the privacy–collaboration–accessibility triangle, none offers an adequate response to all three. These shortcomings highlight the tension between, on the one hand, the value of open universal scientific knowledge and the need to advance medical knowledge as quickly as possible and, on the other hand, the need to protect the privacy of participants and guarantee their full autonomy in participating in research. For these reasons, there is a need to consider alternative approaches. Federated analysis – where the data remains under the control of its initial custodian at all times – is the most promising alternative to managing risks inherent to sharing an individuals genomics data. We previously presented NGS-Logistics as a first approach to federated analysis of NGS-data, offering single-locus regenotyping of federated data hosted at local infrastructure[1]. In this study, we present a more performant strategy, keeping sensitive genotyping data locally and querying out to get restricted and statistical results back [1]. By “bringing computation to the data”, WiNGS offers benefits such as tight control of individual queries (only some queries can be executed, other analyses are not possible), reduced administrative barriers and shared infrastructural requirements. The downside of a federated analysis however, is a more complex technical setup and less flexibility in analysis strategies.

Our Widely Integrated NGS (WiNGS) platform sits at the crossroad of patient privacy rights and the need for highly performant and collaborative genetic variant interpretation platforms. It is a fast, fully interactive, and open source web-based platform to analyze DNA variants in both research and diagnostic settings. Its architecture is developed as a modular and federated setup with a focus on security, privacy-control, and performance.

## 2. Material and Methods

WiNGS was designed as a modular and multi-tier platform, with robustness and flexibility in mind. Its general layout is visualized in Figure 1. Centrally, WiNGS features an ASP.NET and MS-SQL user-interface, referred to as WiNGS-UI, which handles access control, data management, visualization, and all communication with the federated client hubs. Client hubs, referred to as WiNGS-Clients, are deployed on premise as containerized NodeJS/MongoDB applications and manage all sensitive data. Based on this layout, various functional modules, described below, provide means to interact with the data from the central interface.

**Figure 1.**
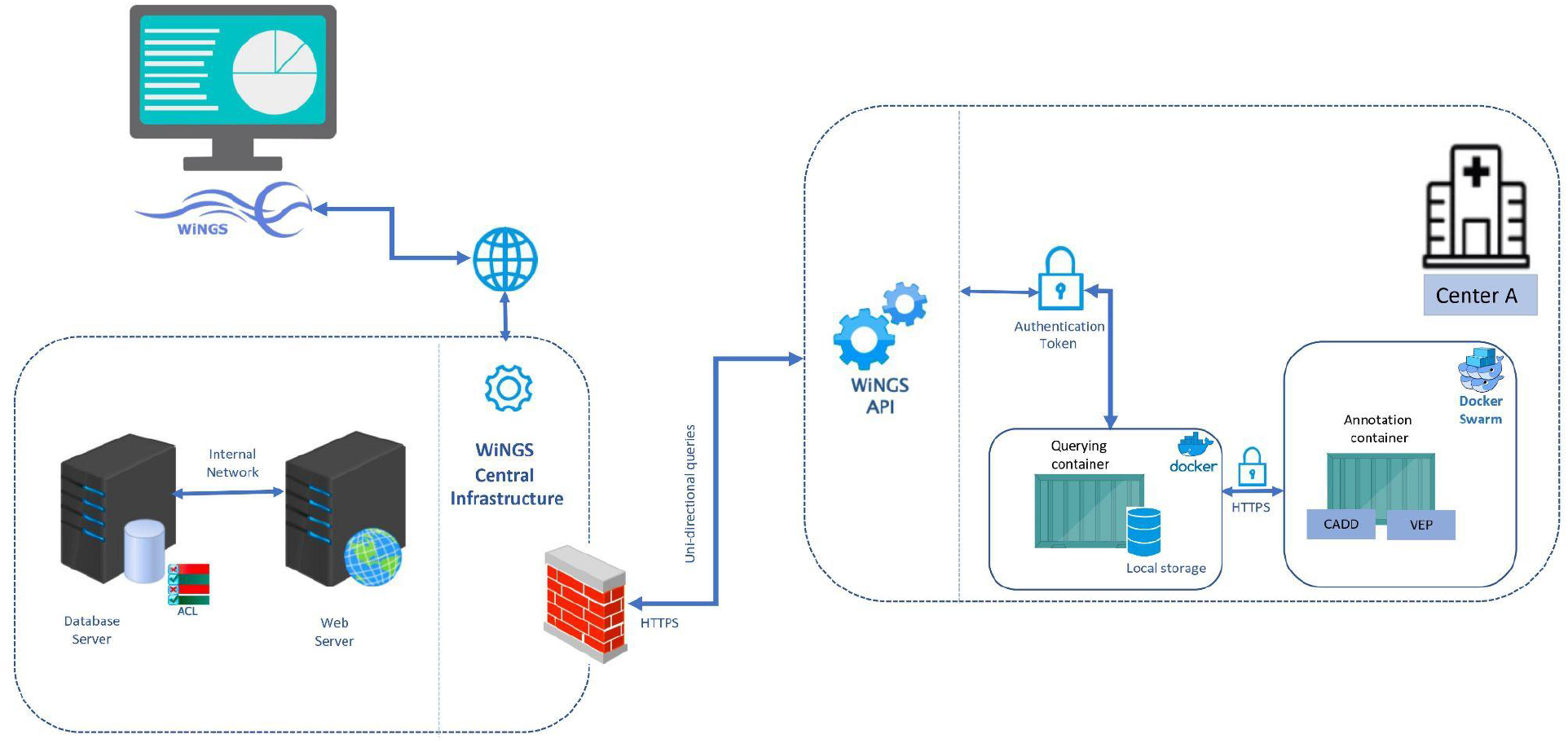
WiNGS components: Central Infrastructure and API. Users log in through the web interface. Access to data is restricted based on the access control list. Queries sent via the user interface are forwarded by the central database server, secured through both https encryption and JSON Web Token authentication, to the docker (client infrastructure) via the internal WiNGS API. Each center has its own database and responds back query results to the central database, for display via UI.

### Client deployment

Within the WiNGS ecosystem, sensitive data is kept on premise, at the client site. This information involves both genetic information (e.g., variant data) and patient information, such as phenotypes, names, or pedigree information. MongoDB, a noSQL solution was selected to store the data because of its greater performance and flexibility [16]. Communication with the central WiNGS server is provided through a NodeJS REST API, secured through both https encryption and JSON Web Token authentication. The REST API handles the queries issued by the user and propagated by the WiNGS-UI, as discussed in the following sections. Hardware requirements are listed in Table 1.

**Table 1:** Minimal hardware requirements for WiNGS client installations.

|  | Storage Module | Annotation Module |
| --- | --- | --- |
| Operating System | Ubuntu 16.04+, kernel 2.6.36+ | Ubuntu 16.04+, kernel 2.6.36+ |
| CPU | 4 | 8 |
| Memory | 32GB, DDR4@3600Mhz | 32GB, DDR4@3600Mhz |
| Storage | 4Gb/WGS sample | 1.6 TB |

In addition to the storage and querying module, WiNGS-Clients can optionally host a full annotation stack. Presence of this stack ensures that variant information for imported samples does not leave the premises. Alternatively, novel variants can be sent to a centralized deployment for annotation. Both options are available to allow clients to balance privacy and hardware requirements, as variant annotation has significant computational requirements with high peak loads.

The annotation stack consists of an Annotation Router and Parser, handling requests and dependent jobs, and local installations of Ensembl VEP and CADD. The Router and Parser have been designed to seamlessly accept additional annotation sources in the future. A Docker swarm is used to enable load balancing and scaling if the local infrastructure permits. Similarly to the storage and querying module, the annotation stack is provided in a containerized format, encapsulating all dependencies and installation routines. Specifically, Ensembl VEP and CADD v1.6 installation scripts have been customized to set up the cache files and data sources for both Genome build hg19 and hg38.

### Access control

With a strong focus on privacy, the access control module of WiNGS, referred to as WiNGS-ACL, forms the core of the platform. Access is regulated in a twofold manner within WiNGS. First, users are organized as illustrated in Figure 2. For each registered client entity, e.g. a hospital, a local administrator is assigned, who can monitor data imports and manage user registration. Second, one or more Principal Investigators are assigned. The minimal setup is to assign each new imported data set to a PI. When a regular user registers in the system, he or she is assigned to a specified PI, which allows this user to access the data associated with the PI. Furthermore, a user can be assigned to a second PI by the Center Administrator (blue dashed line in Figure 2a) to collaborate on the data of that PI. Alternatively, users can be assigned to user groups (green shaded area), to which data can be exposed later (see below).

**Figure 2:**
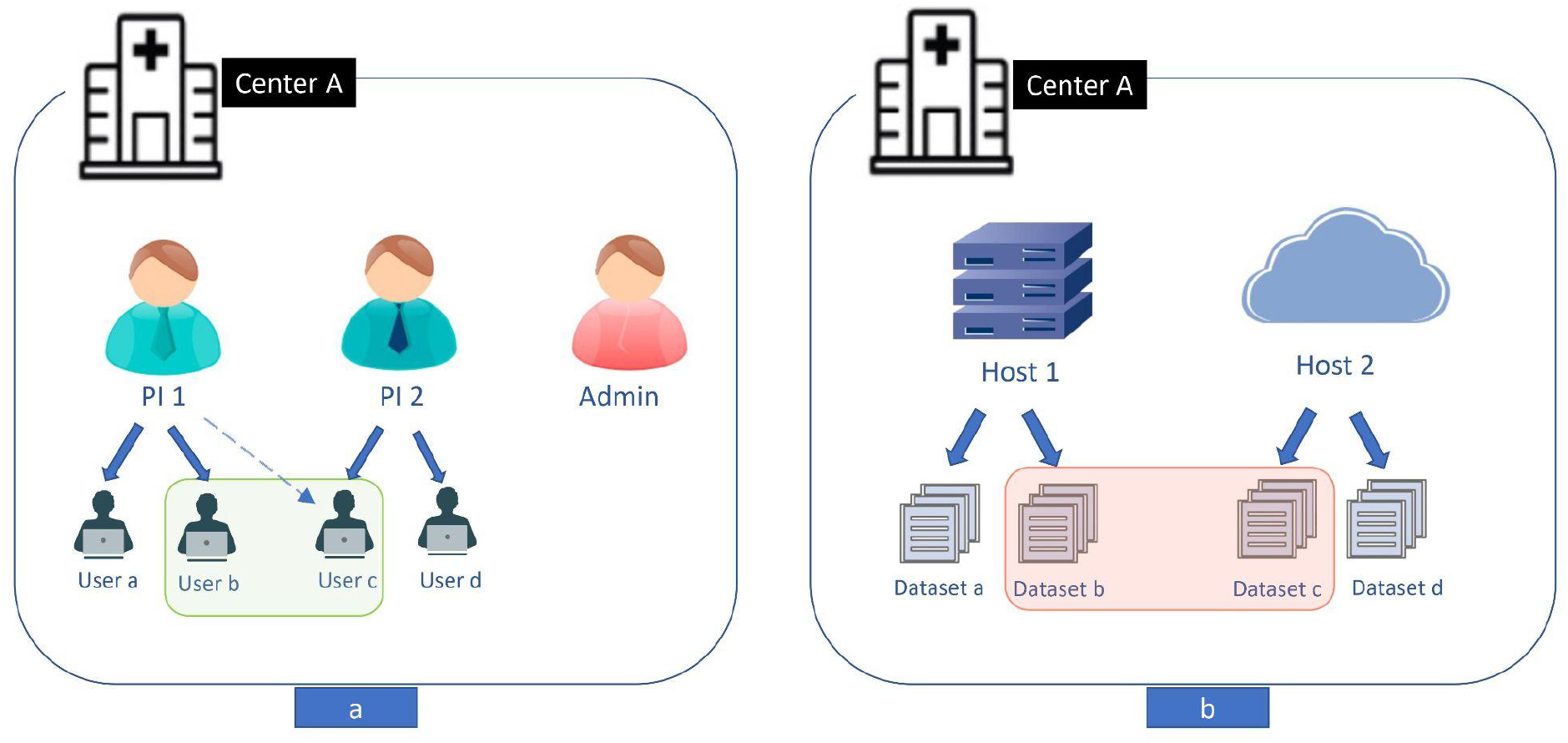
User (a) and data (b) hierarchy layout underlying the WiNGS Access Control List. User hierarchy is defined per client center, restricting user groups (green area) to single centers. Data hierarchy is defined per PI in a center, as illustrated on the right for one PI. Data sets (red area) are thus restricted to data from a single PI in a single center. Access to data sets can also be granted across center users and user groups.

Complementary to the user hierarchy, data is also hierarchically organized, as illustrated in Figure 2b. For each client center, WiNGS supports two physical installations, or hosts, referred to as WiNGS-Clients. For example a hospital can decide to deploy a WiNGS-client both on premise and in the cloud. Each imported data set is physically located in one WiNGS-client and assigned to a single PI. This makes the data available to all users associated with this PI. By default, all data is assigned to a ‘main’ data set, and depending on the specified data type, to a research or diagnostics data set. When data needs to be shared with third parties, custom subsets (red area in Figure 2b) of data can be created and shared by the PI. Note that potentially required processes, such as ethical approval, are outside the scope of WiNGS and left to the responsibility of the PI. Access can be granted to both user groups and individual users across the whole WiNGS ecosystem. To monitor data access, PIs can request the list of users and the data sets to which they have access through the user interface.

### Data import

The base data entity within WiNGs is the individual (see Figure 3). All information and data related to an individual, including data about family members, is stored in a single WiNGS-client to make precomputation possible (see below). The minimal information requirement for individuals is sex (female/male/unspecified), and further registration is based on a pseudonymized identifier. Support for identifiable metadata, such as birth date and names is available, although users should keep data minimization principles in mind. Next, one or more experimental data sets, reflecting the type of sequencing (whole genome, whole exome, or panel resequencing) can be linked to the individual, referenced within WiNGS as ‘Samples’. Finally, each Sample can be associated to one or more sets of ‘Files’, reflecting data analysis results in a specific genome build. Variant information can be imported from both VCF and gVCF files. BAM, CRAM and FASTQ files can also be managed in WiNGS.

**Figure 3:**
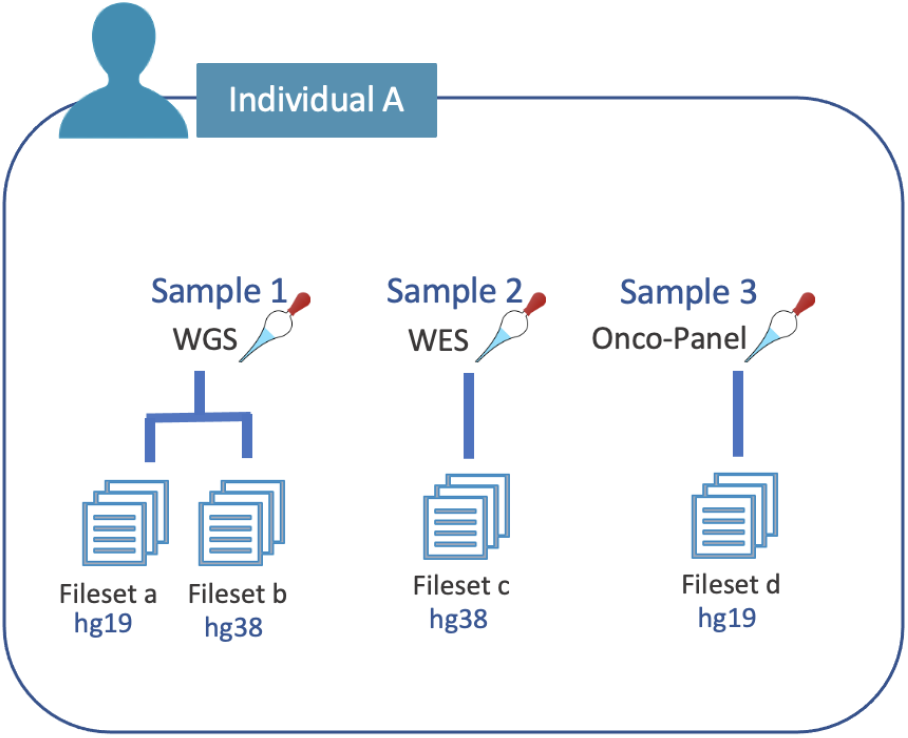
Structural links between individuals and data sets.

**Figure 4:**
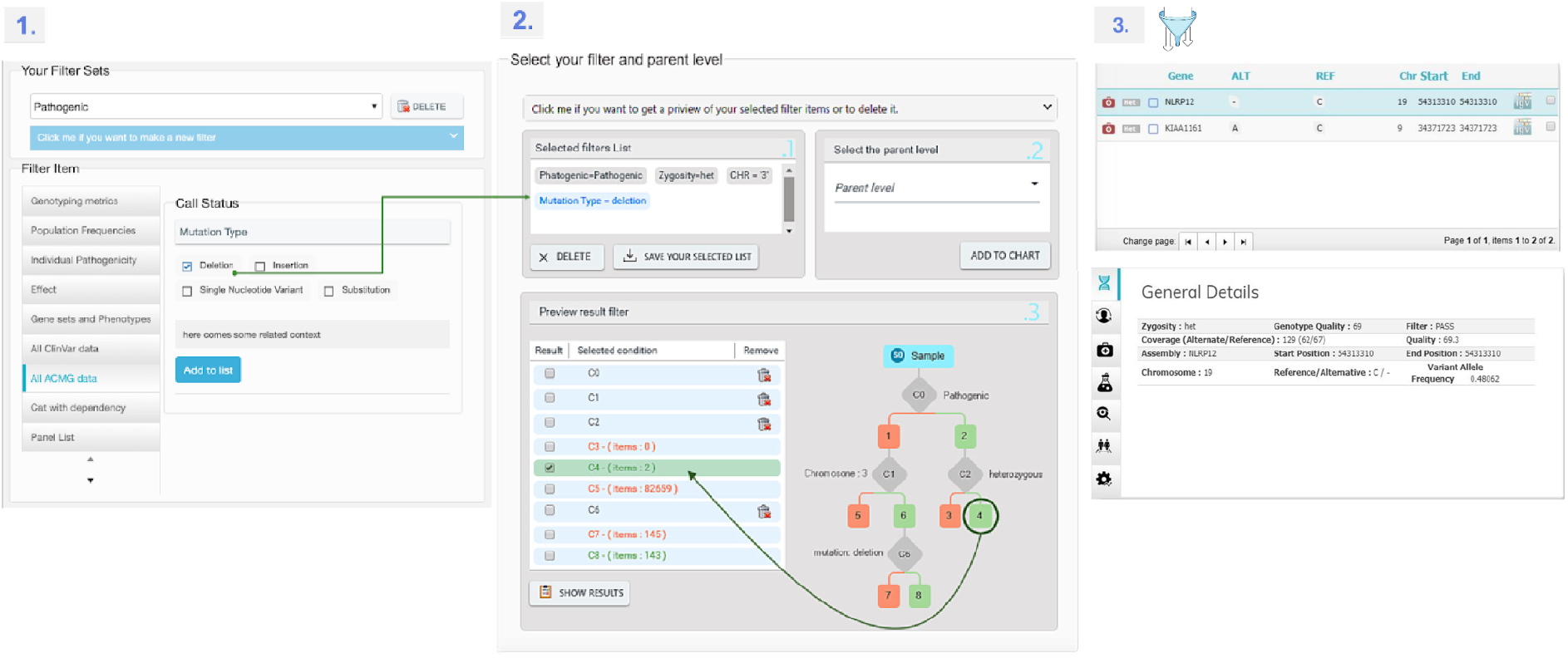
Filtering interface. Users select annotation criteria to apply as filter (Panel 1), and add them to the filtering tree (Panel 2). After selection of relevant leafs, passing variants and annotations are shown in the results page (Panel 3).

There are two approaches to import data into WiNGS, either interactively through the WiNGS-UI or via an automated procedure based on a sample sheet. The latter process is aimed at facilitating and automating sample import. Sample sheets, CSV-based files located at the WiNGS-Client, are queried daily by the central system to fetch new samples for import. Individuals, samples, and data sets are automatically created and linked, followed by import of the (g)VCF file at the WiNGS-Client. After completion, the center administrator is notified through an email including general results and potential issues. Additionally, full procedure logs are available to the admin through the WiNGS-UI. During manual import, individuals and samples should be registered prior to adding file(s). To add the files, a free text box is provided to get the path of the file from the user, supporting both paths at the WiNGS-Client, or URLs.

### Variant annotation

After importing a (g)VCF file, variants are sent for annotation. A two-tier approach was implemented to improve performance. First, annotations from variants seen before at the importing WiNGS-Client are recycled and added to the newly imported variants. Second, the remaining, novel variants are sent to the annotation stack for processing, significantly reducing the computational requirements. Variants are annotated using ensembl-vep:release_105.0 with gene information based on RefSeq transcripts, population frequencies from gnomAD and disease associations available in ClinVar. Finally, deleteriousness predictions are obtained using CADD v1.6. On successful import and annotation, the sample is marked as available for analysis.

### Variant querying

WiNGS features a powerful and versatile module for a variety of analyses, including single sample, trio-based, and variant-based approaches. To ensure privacy, users can perform analysis only on the samples to which they have been granted access. The WiNGS-ACL translates requests into queries forwarded to specific WiNGS-Clients, based on the users access rights. These queries contain the criteria to be applied on the variant annotations, such as variant coordinates and genotype, impact and pathogenicity predictions, population frequencies, family inheritance patterns and clinical annotations. Criteria can be combined using a graphical tree-structure, providing real-time feedback of variants passing the specified criteria. The system administrator has access to a module in the WiNGS-UI where mongoDB document fields are translated into UI filtering elements. This way, future annotation extensions, for example a novel pathogenicity score, can easily be made available to users as a filtering criterion.

In addition, specific strategies were put in place to improve performance and maintain data privacy based on the analysis approach. For trio analysis, WiNGS leverages predefined pedigree info and precalculated segregation statistics to improve querying efficiency. Once the precomputing phase is complete for a trio, the user can proceed to analyze a subset of interest, for example *de novo* variants, using the various functional filters common to all approaches.

Despite the obvious security value of keeping sensitive data at the clients infrastructure, it is mainly with variant-based querying that the federated analytics module of WiNGS shows its benefits. Queries based on a variant, or gene, are sent to all client instances in an asynchronous routine as a minimal request for allelic counts. Automatic translation between genome builds allow for seamless interrogation of the full dataspace of WiNGS. Upon data retrieval, variant counts are aggregated by the WiNGS-UI. Ultimately, variant statistics are reported containing only public information, similar to what is provided by the Beacon framework. Details presented here include systemwide counts per variant, the effect of the variant on the affected gene (if any), and population frequencies (gnomAD).

### Phenotyping

WiNGS supports both structured and unstructured phenotyping of individuals. First, the Human Phenotype Ontology (HPO) is available as a source for the annotation of structured phenotypes. Second, the Online Mendelian Inheritance in Man (OMIM) database is embedded in WiNGS to associate HPO terms to related diseases. Finally, genes known to be associated with selected HPO terms, data is taken from HPO community, can be extracted from the provided phenotypes, or be provided as a source of phenotypes by the user. To facilitate HPO-term selection, the graphical path around HPO terms of interest is provided as part of the phenotyping module. As an extension to binary phenotyping, the age-of-onset and severity of each HPO-term can be provided. Alternatively, users can enter free text to describe the clinical phenotype.

## 4. Results

In the pilot phase, WiNGS is deployed over four centers with one hosts each. The total number of imported WGS samples is 516 over all centers. We evaluated the performance of WiNGS for different analyses such as variant discovery, single sample, and trio analysis. Variant discovery analysis was evaluated among four registered centers, 3 trio cases were evaluated in 2 different centers, and sample discovery was evaluated for 15 samples.

### Performance

Several performance aspects were evaluated. The first, and typically most time consuming step in variant interpretation is data annotation. Within WiNGS, this step is performed by the WiNGS-Client infrastructure. Import consists of two steps, namely data g(VCF) parsing and annotation of novel variants. Figure 5 demonstrates that after import of a limited number of samples, most common variants are present in the system, after which import times are reduced to approximately two hours per genome. Because the import phase is not affected by the total size of the database, it can be expected that over time, the total import time will further reduce to the baseline of 30 minutes for a genome sample.

**Figure 5:**
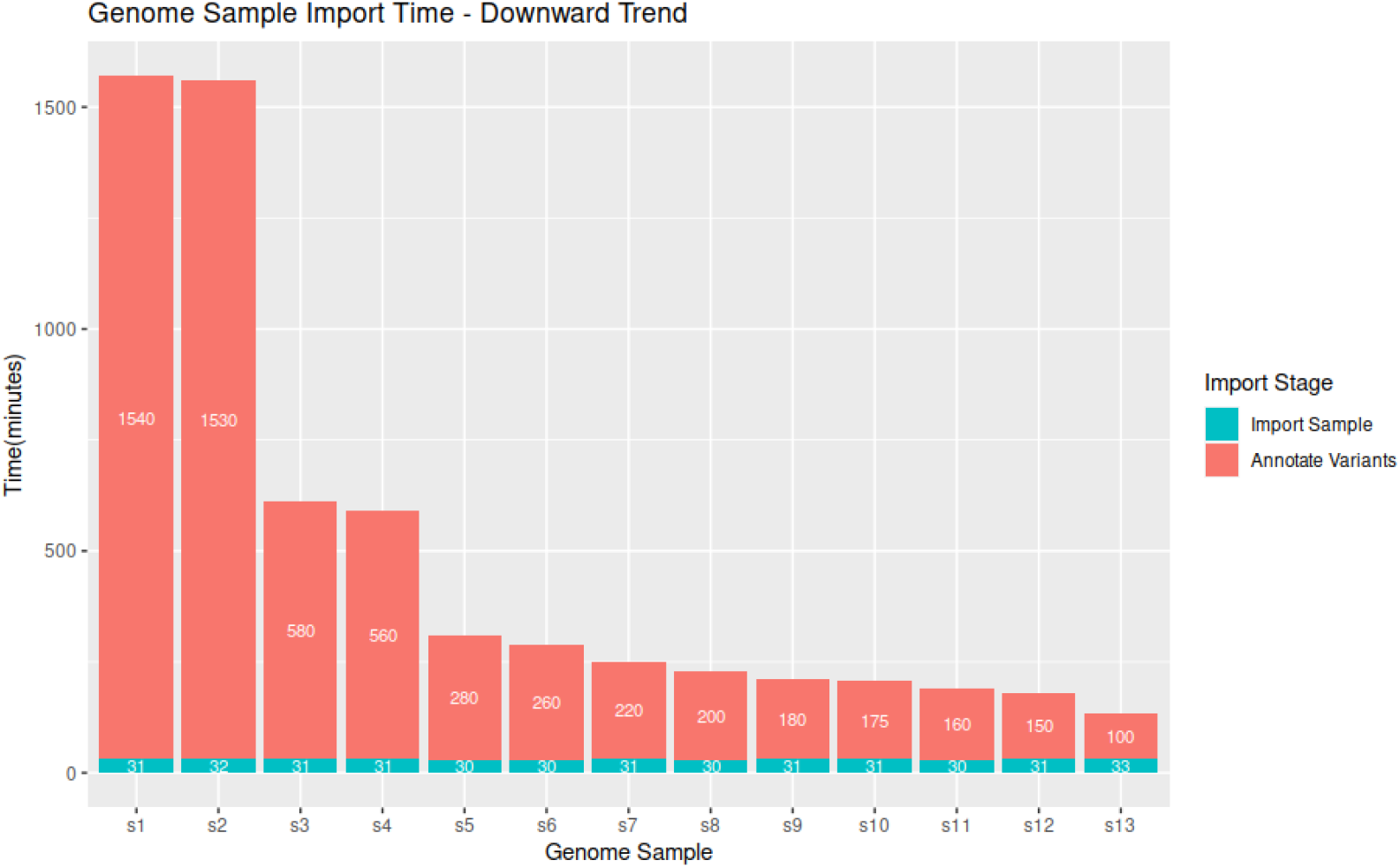
Sample import time decreases with the number of novel variants that need to be annotated.

After data import, which can be automated as a nightly process through samplesheets, the most significant computational task is precomputation of variant segregation (see Table 2). This one-time action is performed automatically when individuals are combined into families through the pedigree module.

**Table 2:**
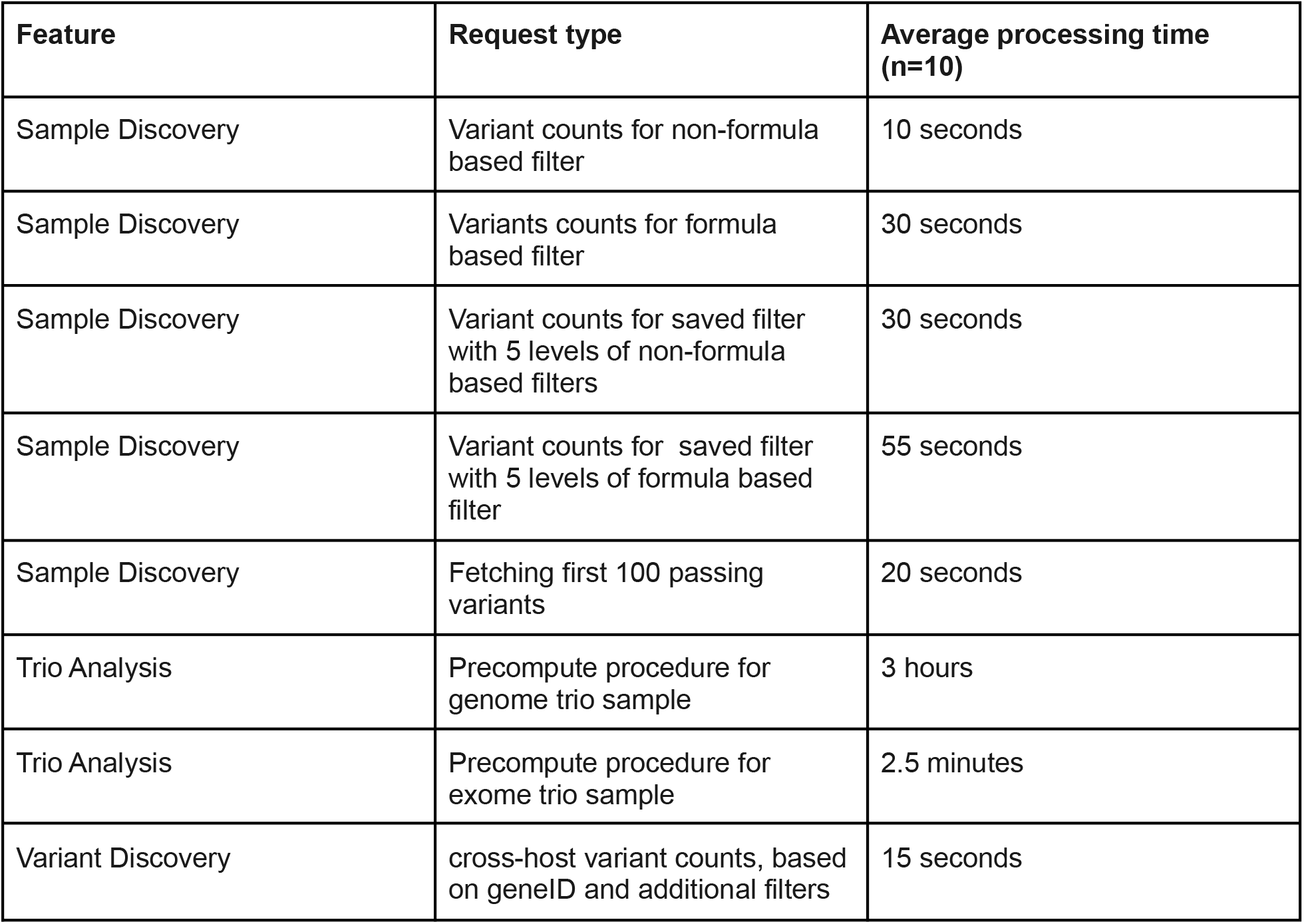
Performance Metrics of the WiNGS platform. Query performance was averaged over 10 executions per scenario. Variants counts: variants passing added filter criterium. Saved filter: Re-application of a filter tree on a novel sample. Fetch passing variant: Retrieve variants and annotations on results page. Non-formula : direct query on annotation values. Formula : query on combined annotation values (e.g. sum).

Finally, actual querying was benchmarked using a center with 400 genomes samples. Multiple centers with different hardware and genome samples were used for the *Variant Discovery* tests. Using *Sample Discovery*, two different filter types were evaluated. Non-formula based filters used in these tests directly query stored data fields, including gnomAD allelic frequency, ClinVar, Variant type and chromosomal region. Formula based filter types require computation on multiple stored values, and include total depth or allelic ratio. Both filter types were evaluated for three distinct request types, namely the number of variants passing newly set filters, the number of variants passing a previously saved combination of filters, and fetching actual results from the client. Results are shown in Table 2.

### Use case

#### Sample Discovery

A synthetic case was constructed based on the presentation of a patient with Intellectual Disability sent in for whole exome sequencing. Because a *de novo* genetic cause was suspected, both parents were also sequenced. The variant identified in our clinical WES setting was spiked into the data of the Ashkenazim Jewish NIST/GiaB Trio (index:HG002, father:HG003:, mother:HG004), downloaded from ftp://ftp-trace.ncbi.nlm.nih.gov/giab/ftp/data/AshkenazimTrio/ [23]. The data was generated using Illumina paired-end sequencing, and analyzed using BWA/GATK in genome build hg19, as described in the publication. The variants were added to the final VCF files. After WiNGS import, a pedigree was constructed for the trio, followed by segregation precomputation. A filtering strategy was constructed based on Quality (VQSR Tranche = ‘PASS’) and the expected absence of the variant in control populations (gnomAD.Allele Frequency < 0.01). Functional impact of the causative variant (VEP.impact = ‘HIGH’) was added to the criteria, as well as the complementary CADD pathogenicity prediction (CADD.phred > 25) (figure 6).

**Figure 6:**
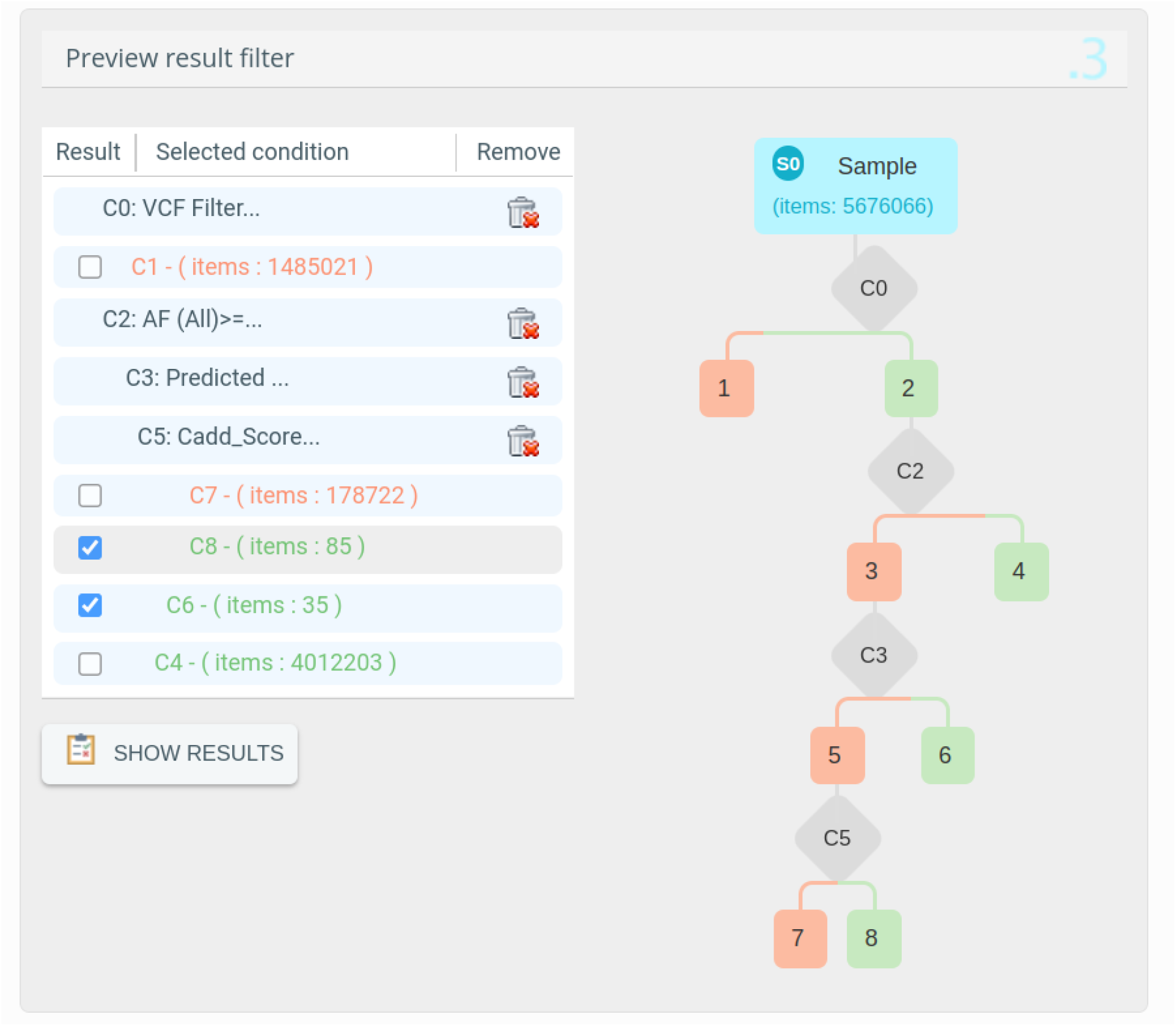
Sample Discovery preview of the passing variants. Using four criteria, the amount of candidate variants is reduced to less than 0.05% of the variants present in the index patient.

Without looking at family information, 120 out of 5.6 million variaints passed these criteria. It should be noted here, that high CADD.phred scores (node C6) do not necessarily reflect a “HIGH” predicted functional impact. Hence, we would select both leaves in the tree for further inspection. Although the resulting variant set could be reduced using other available criteria (e.g. known phenotypic association of the affected genes), we can apply the expectation of a *de novo* inheritance, by switching to the “*Trio Analysis”* module, offering an intuitive selection of specific segregation patterns (figure 7), 5 variants were retained under the hypothesis of a *de novo* mutation. Further analysis on the allele frequency across all registered centers against the patient phenotype, identified a second carrier of one of a variant in PUF60, presenting with Intellectual Disability, and no unaffected carriers (see figure 9). This results in a significant correlation between the phenotype “Intellectual Disability” (HPO:0001249) and the variant NM_001271096.2:c.404G>T_p.Gly135Val (Fisher Exact < 0.01). Literature review showed that disruptive variants in *PUF60* are associated with Verheij Syndrome, which is in line with the phenotype of the presented case. The remaining variants either showed not additional affected carriers, or showed unaffected carriers, resulting in non-significant correlation.

**Figure 7:**
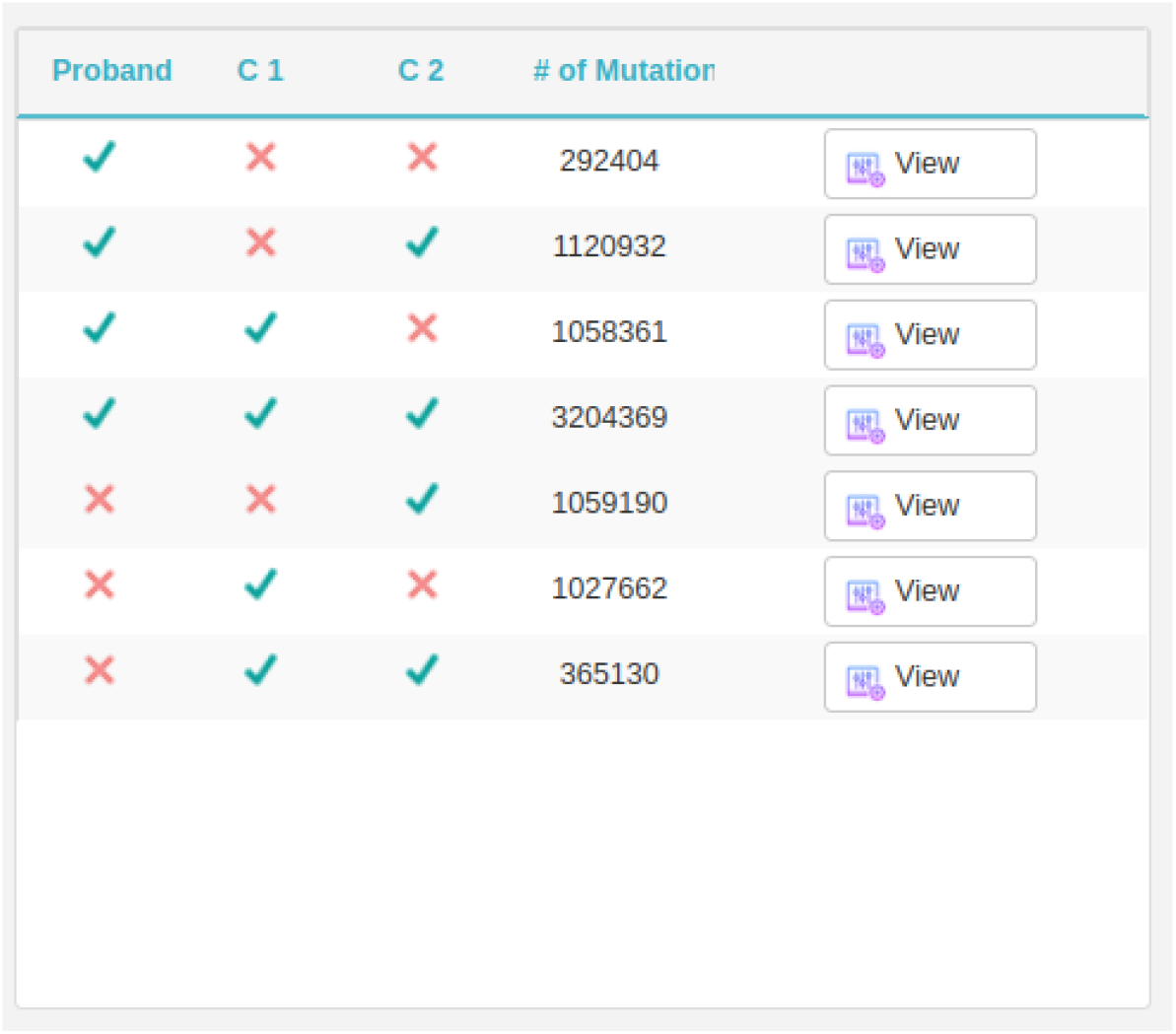
Overview of precomputed variant segregation results. Numbers reflect the amount of variants seen or absent in specific sample combinations, taking into account both VCF and gVCF reference calls.

**Figure 8:**
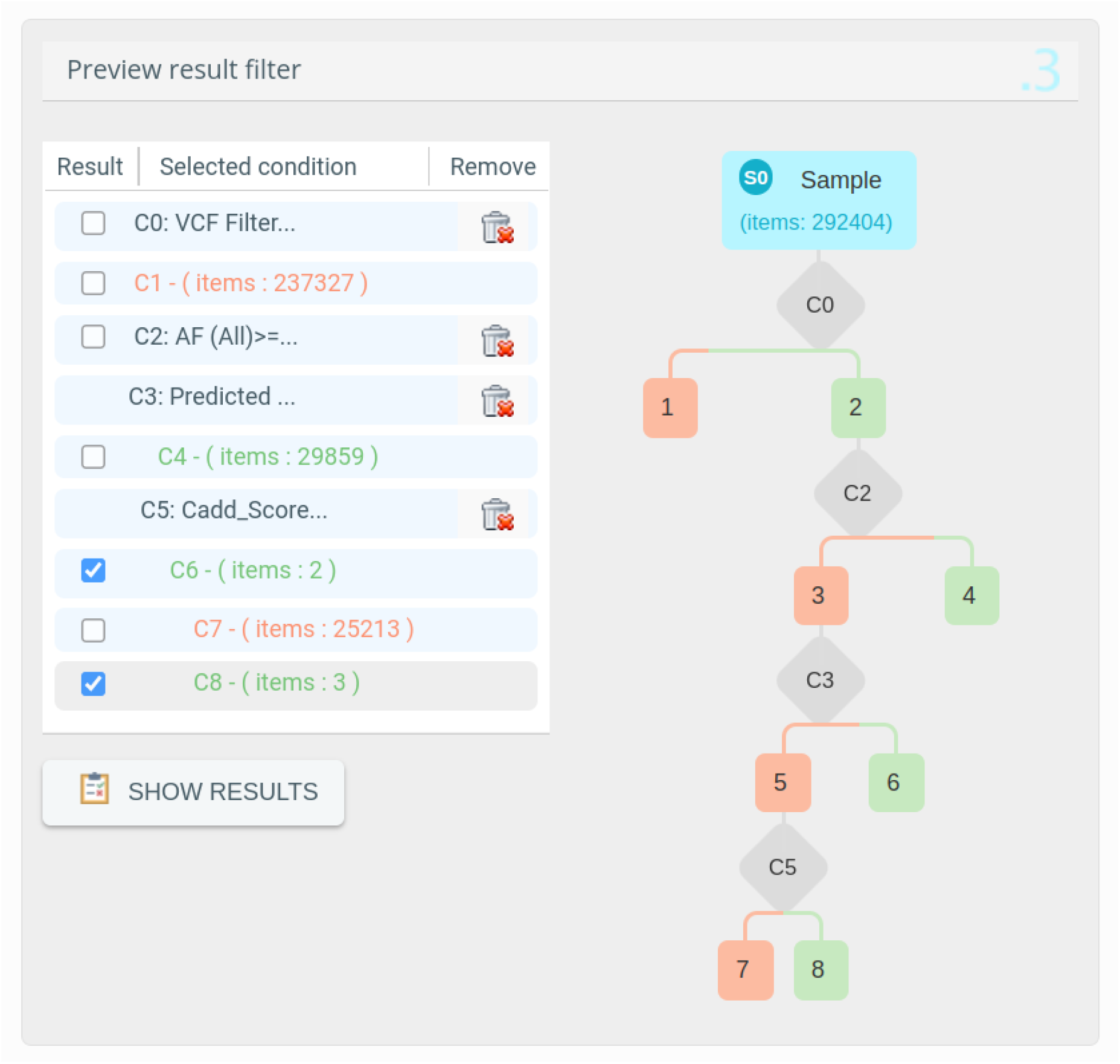
Trio Analysis preview of the passing variants under the de novo hypothesis. Using the segregation information, 5 variants pass the criteria from Figure 6.

**Figure 9:**
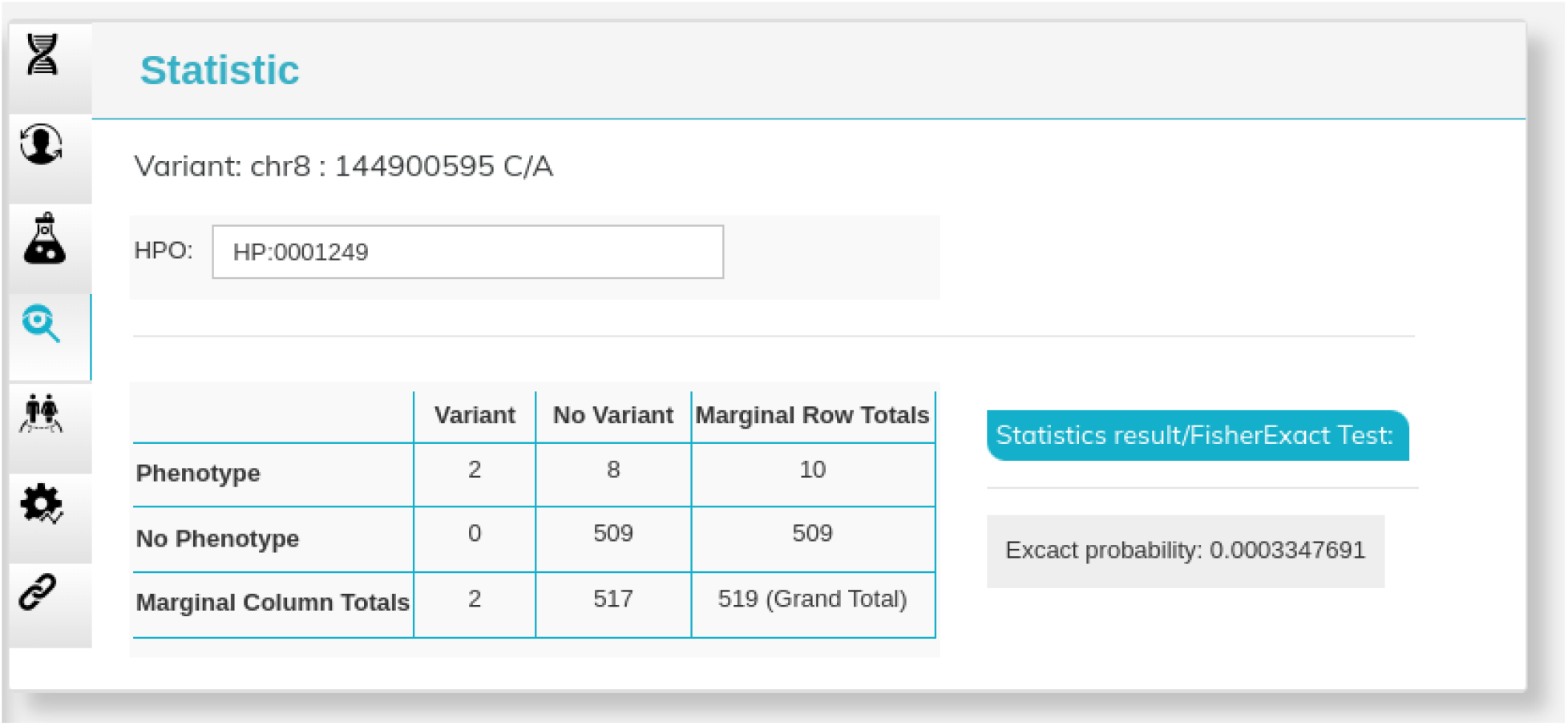
Cross-client genotype/phenotype association testing

## 5. Discussion

WiNGS has currently matured to a strong proof-of-principle platform for federated interpretation and analysis of genomic variants data. It fills an important gap in solutions for privacy-preserving personal genomic data sharing with an innovative federated analytics approach that smoothly dovetails with GDPR requirements regarding data minimization. We address several of the currently standing challenges in the analysis of large scale genomics data, including data security, accessibility and interpretability, by moving from a centralized to a federated model.

Centralized data hubs such as dbGaP and EGA, hosting raw and processed data, ship part of this data across multiple research or clinical centers for reanalysis purposes. Although, they provide a tight initial level of control, through verification of the credentials of researchers, compatibility of research goals with consent and ethical approvals, the legitimate bureaucratic barriers created by centralized controlled-access databases are an important hurdle that discourages some potential users from completing the cumbersome approval processes, which results in underuse of the data. As an alternative, platforms can offer ‘federated analytics’ solutions, by which we mean an analytics infrastructure where personal data (here, sensitive genetic and clinical data) does not leave the infrastructure of its original data controller. Rather than “sending out the data for analysis”, “the analysis is sent to the data”. Within WiNGS, the analysis currently consists of specific, preconfigured data queries, although more complex analyses can be easily enabled in future work. Managing the allowed analytics centrally functions as a safeguard against malicious code deployment.

A second concern is the misuse of data, once redistributed, for unapproved research or non-research purposes (e.g., law enforcement). Pseudonymized data may not always fully prevent re-identification of patients, even without access to the pseudonymization registry. The architectural setup of federated platforms, such as the one presented here, naturally facilitates privacy-by-design and prevents the misuse of redistributed data, as sensitive data is kept on-premise at all times. Furthermore, most federated approaches such as WiNGS or the Beacon project, do not expose full data, but limit access to aggregated statistics. Nevertheless, sensitive data can potentially be obtained from these systems through reconstruction and tracing attacks [17]. Reconstruction attacks happen when sensitive data is revealed about most individuals covered by the dataset. It would happen if from the aggregate statistical data it was in fact possible to reconstruct the data of the underlying individuals. Tracing attacks are a modest adversarial goal referring to the ability to infer whether or not a specific individual is part of the data set [17]. In order to perform a tracing attack, the adversary requires access to the genetic profile of the individual [18, 20, 19], but could for example disclose some medical status (e.g., a particular diagnosis, based on the metadata of the attacked data source). Mitigation strategies against re-identification attacks are possible, such as statistical noise injection for differential privacy [18] or query monitoring to limit systematic query scans. Although these strategies can be implemented in WiNGS, the system can only be accessed by registered users, always affiliated to, and under the authority of a participating center.

Recently, a set of specific privacy guidelines were issued relating to the processing of personal data of European citizens, termed the General Data Protection Regulation (GDPR). Adoption of and compliance with these guidelines is a prerequisite for any analytical platform, to be usable in a European clinical setting. In general, the GDPR provides a tight set of constraints guiding the design of analytics systems to be mindful of privacy, and to likely meet privacy requirements and best practices anywhere in the world. Moreover, as the GDPR applies to all processing carried out both by organizations operating within the EU and to organizations outside the EU offering goods or services to individuals in the EU, it is a key constraint for globally operating entities, including academic consortia.

A key GDPR principle is data minimization, and it is specifically emphasized to choose an architecture with strong data minimization (e.g., federated analytics) over one with poorer data minimization (e.g., centralized databases) if both can adequately fulfill the necessary purposes (https://gdpr-info.eu) *Art. 89(1)*. When considering our federated model (Figure 1), it can be seen that data minimization is central to our design, with tight control of what data is accessible and can be visualized through the UI. With regard to federated analytics, the key idea behind WiNGS is to process the data in such a way that data shared by centers and aggregated by the hub is *not* personal data, but only aggregate statistical data. Specifically, queries sent out to the clients in the *Variant Discovery* module result in aggregated counts, which are sent back to the central infrastructure, where they are aggregated again over all clients. This approach allows us to leverage the GDPR statistical exemption (https://gdpr-info.eu) Art 9(2)(j), *89(1), Recital 162*. Another route to achieve a similar result could be to leverage the closely related, and more familiar, research exemption. However, the statistical exemption is also applicable to the processing of clinical data in routine clinical care, outside the research setting. Hence, WiNGS can leverage cross-institutional frequency metrics as aggregated statistics to perform for example genotype/phenotype associations. Agreements for more general collaborations, requiring different lawful bases and ethical agreements, are left to the users to set up. Once set up, WiNGS then provides the means to grant access to the relevant datasets.

The federated analytics framework of WiNGS can be extended with more advanced methods in future work, as long as the details of the GDPR statistical exemption are kept in mind. The GDPR provides a fairly clear standard of ‘aggregate statistical data’. Data should be aggregated in such a way that it is not personal data anymore, which means in particular that it should not be possible to reconstruct personal data from the available aggregate statistical data. As discussed before, this raises the question of possible re-identification attacks. Second, the definition of ‘statistical purpose’ is less clear as it relies on the notion of ‘statistical result’, which is left undefined. For lack of further guidance, we will consider a ‘statistical result’ as a piece of information relevant to a population rather than to a specific individual. A mean, a standard deviation, a logistic regression model, a deep learning model are ‘statistical results’. The individual data of a pseudonymous patient or the risk prediction score from a logistic model for a specific patient are *not* ‘statistical results’. This interpretation is supported by the caveat that “this result or the personal data are not used in support of measures or decisions regarding any particular natural person”. Hence, we can leverage all data within the WiNGS platform to train classification models for disease-related variants, and offer them to the users for application on their respective accessible data.

A final benefit of federated frameworks, is that although large centralized databases can be better shielded against third-party attacks than privately hosted databases, they create a single point of failure and can be opaque to government interference such as search warrants or national security interventions (https://en.wikipedia.org/wiki/Room_641A). While some research data in the US might be protected from search warrants via certificates of confidentiality [15], the situation is less clear in Europe as law enforcement and intelligence activities do not fall under the GDPR, and regulations in China foresee broad access to any genetic resource by authorities for public health, national security, and societal public interest purposes (https://www.chinalawtranslate.com/en/p-r-c-regulation-on-the-management-of-human-genetic-resources/) Art 16. As all sensitive data within our federated framework is kept at the data controller’s premises, data security is handled at the institutional level. On top, the central WiNGS hub has no routines to extract all data from the client’s hub, preventing privacy breaches by an attack or warrant on the central infrastructure.

Other federated frameworks exist in the context of genomics, including DataShield [22]. However these platforms have a more low-level analysis focus, aimed at raw data analysis rather than variant interpretation. Although these platforms could be used to query federated VCF data using command-line tools, the high performance of the WiNGS noSQL backend allows us to perform these tasks in realtime using a more intuitive web-interface. In contrast, several web based platforms are available to facilitate variant interpretation. Two examples are Diploid’s Moon platform (https://www.diploid.com/moon) and Genoox’s Franklin platform (https://www.genoox.com/). Although widely used, these platforms are examples of centralized applications requiring upload of sensitive information, which can, depending on the institutional data protection guidelines, be prohibitive to their implementation. Several extensions to the WiNGS platform can be envisioned to further bridge the gap between existing efforts, while maintaining stringent data privacy guidelines. First, a higher granularity of access control might be based on dataset, instead of Principal Investigator, allowing to assign data from different projects under a single PI, to different users. Second, automatization of the platform through a RESTful API can integrate the platform with existing efforts such as the GA4GH Beacon project or private analysis pipelines. Finally, the federated analytics system can be applied to train variant classification models.

## 6. Availability

The web server can be accessed via the following link: https://wings-platform.org/ The source code is available under the Affero general public license version 3, at the following location: https://github.com/wings-public

A demonstration of the platform is available at https://www.youtube.com/watch?v=wLviKPVgoSQ

Data from the presented use case is available from the WiNGS system as a public data set, named “Demo Data”

## 7. Acknowledgement

H Ch, N Sh, YM are also funded by Research Council KU Leuven: Symbiosis3 (C14/18/092); Federated cloud-based Artificial Intelligence-driven platform for liquid biopsy analyses (C3/20/100); CELSA - Active Learning (CELSA/21/019), CELSA-HIDUCTION (CELSA/17/032), Flemish Governmen (FWO: SBO (S003422N), Elixir Belgium (I002819N); SB and Postdoctoral grants, AI Research Program, VLAIO PM: Augmenting Therapeutic Effectiveness through Novel Analytics (HBC.2019.2528)), EU (This project has received funding from the European Union’s Horizon 2020 research and innovation programme under the Marie Skłodowska-Curie grant agreement No. 956832.)

